# Loss of PI3-kinase activity of inositol polyphosphate multikinase impairs PDK1-mediated AKT activation, cell migration and intestinal homeostasis

**DOI:** 10.1101/2020.12.18.423145

**Authors:** Prasun Guha, Luke Reilly, Evan R. Semenza, Efrat Abramson, Subrata Mishra, Yoshitasu Sei, Stephen A. Wank, Mark Donowitz, Solomon H. Snyder

**Affiliations:** The Solomon H. Snyder Department of Neuroscience, Johns Hopkins University School of Medicine, Baltimore, MD 21205, USA; Department of Neurology, Johns Hopkins University School of Medicine, Baltimore, Maryland 21205, USA; Digestive Diseases Branch, National Institute of Diabetes and Digestive and Kidney Diseases, National Institutes of Health, Bethesda, Maryland; Department of Medicine, Division of Gastroenterology, Johns Hopkins University School of Medicine, 720 Rutland Avenue, Baltimore, MD 21205, USA; Department of Psychiatry and Behavioral Sciences, Johns Hopkins University School of Medicine, Baltimore, MD 21205, USA; Department of Pharmacology and Molecular Sciences, Johns Hopkins University School of Medicine, Baltimore, MD 21205, USA; Department of Biophysics and Biophysical Chemistry, Johns Hopkins University School of Medicine, Baltimore, MD 21205, USA; Reference Standard Laboratory, United States Pharmacopeial Convention, Rockville, MD, 20852, USA

**Keywords:** Crohn’s disease, Inflammation, IPMK, AKT, PDK1, Inositol, PI3kinase, Intestine, Ileum, Regeneration, Paneth Cell, Chemoprotection

## Abstract

Inositol polyphosphate multikinase (IPMK) is a rate-limiting enzyme in the inositol phosphate (IP) pathway which converts IP3 to IP4 and IP5. In mammalian cells, IPMK can also act as a phosphoinositol-3-kinase (PI3-kinase). We previously found that IPMK is a critical PI3-kinase activator of AKT. Here, we show that IPMK mediates AKT activation by promoting membrane localization and activation of PDK1. The PI3-kinase activity of IPMK is dispensable for membrane localization of AKT, which is entirely controlled by classical PI3-kinase (p110*α*,ß, *γ, δ*). By contrast, we found that PDK1 membrane localization was largely independent of classical PI3-kinase. Membrane localization of PDK1 stimulates cell migration by dissociating ROCK1 from inhibitory binding to RhoE and promoting ROCK1-mediated myosin light chain (MLC) phosphorylation. Deletion of IPMK impairs cell migration associated with the abolition of PDK1-mediated ROCK1 disinhibition and subsequent MLC phosphorylation. To investigate the physiological relevance of IPMK-mediated AKT activation, we generated mice selectively lacking IPMK in epithelial cells of the intestine, where IPMK is highly expressed. Deletion of IPMK in intestinal epithelial cells markedly reduced AKT phosphorylation and diminished numbers of Paneth cells – a crypt-resident epithelial cell type that generates the physiological niche for intestinal stem cells. Ablation of IPMK impaired intestinal epithelial cell regeneration basally and after; chemotherapy-induced damage, suggesting a broad role for IPMK in the activation of AKT and intestinal tissue regeneration. In summary, the PI3-kinase activity of IPMK promotes membrane localization of PDK1, a critical kinase whereby AKT maintains intestinal homeostasis.

**One Sentence Summary:** PI3-kinase activity of IPMK is essential for activation of AKT.

## Introduction

Inositol polyphosphates are signaling molecules generated by an evolutionarily conserved family of kinases. Inositol 1,4,5-trisphosphate (IP3) is well established as a physiologic regulator of intracellular calcium ^1^. Among the family of kinase enzymes regulating inositol phosphates, IPMK is notable for its generation of multiple inositol phosphates, including IP4 and IP5 ^2^. Additionally, IPMK serves as a phosphoinositide 3-kinase (PI3-kinase) in mammalian cells ^3^. The cytoplasmic PI3-kinase activity of IPMK, like that of classical PI3-kinase enzymes, contributes to AKT activation ^3^. Independent of its kinase activity, IPMK mediates autophagy by binding to AMP-dependent protein kinase and the autophagy-activating kinase ULK1 ^4^. IPMK also participates in nutrient sensing and activation of mTORC1 and mTORC2 ^5^. Epidemiological studies indicate that mutation of IPMK at its nuclear localization signal is associated with familial intestinal carcinoids ^6^. Genome-wide association study (GWAS) data from patients with inflammatory bowel disease (IBD) have established IPMK as a putative risk gene ^7^.

AKT is a central player in the regulation of metabolism, cell survival, motility, transcription, and cell-cycle progression ^8^. Classical PI3-kinase (p110*α*,ß,*γ, δ*) generates the second messenger PIP3 from PIP2. This stimulates the translocation of AKT from the cytoplasm to the plasma membrane, which involves its pleckstrin homology (PH) domain. The serine/threonine kinase phosphoinositide-dependent kinase 1 (PDK1) also contains a PH domain and is recruited to the plasma membrane by PIP3. PDK1 phosphorylates AKT at Thr308 ^8^, while phosphorylation at Ser473 is mediated by mTORC2 ^9^. Phosphorylation of AKT at both Thr308 and Ser473 generates an active form of AKT, which targets specific proteins in the cytoplasm and nucleus. Several studies imply that classical PI3-kinase is not the critical enzyme for recruitment of PDK1 and mTORC2 to the membrane to activate AKT ^10, 11, 12^. PIP3-mediated PDK1 membrane localization is also important for activation of Rho-associated coiled-coil-containing kinase 1 (ROCK1), an essential mediator of cell migration. PDK1-mediated ROCK1 activation appears to be only marginally influenced by inhibition of classical PI3kinase ^12, 13^, suggesting a different PI3-kinase mediates this effect. Deletion of IPMK abolishes activation of AKT ^3^ via heretofore obscure mechanisms. Here we demonstrate that IPMK is the principal PI3-kinase regulating PDK1’s membrane localization as well as associated AKT activation and ROCK1-mediated cell migration. We further show that IPMK physiologically influences self-renewal of the intestinal epithelium.

## Results

### IPMK mediates AKT activation

To confirm a role for IPMK in AKT activation, we used wild-type (WT) and *Ipmk* knockout (KO) mouse embryonic fibroblast (MEF) cells developed in our lab ^3^. Cells were serum-starved overnight, followed by treatment with serum for 5 minutes to evaluate AKT phosphorylation. *Ipmk* deletion markedly diminished AKT phosphorylation at Thr308 (Figure 1A, B). Loss of IPMK also abolished AKT phosphorylation at Ser473 (Supplementary Figure 1A). In agreement with previous findings, ^14 8^ we found that deletion of *Ipmk* impaired mTORC2 kinase activity and thus reduced AKT Ser473 phosphorylation (Supplementary Figure 1B). We explored mechanisms underlying deficits in AKT Thr308 phosphorylation in *Ipmk*-deleted cells. AKT phosphorylates GSK3β at Ser9 to inactivate GSK3β ^8^. IPMK KO MEFs showed a substantial loss of GSK3β phosphorylation at Ser9 (Figure 1A, C). To determine whether phosphorylation of other AKT substrates might be defective in IPMK KO cells, we used an antibody recognizing proteins containing phosphorylated AKT substrate domains. Phosphorylation detected by this antibody was considerably reduced in IPMK KO MEFs and was comparable to that of cells treated with the pan-PI3-kinase inhibitor pictilisib (Figure 1 D). A time-course study showed that deletion of *Ipmk* substantially reduced insulin-stimulated AKT Thr308 phosphorylation (Figure 1 E, F). We extended our finding to another cell type. Knockdown of *IPMK* by shRNA in HEK293 cells (Supplementary Figure 1C) substantially reduced phosphorylation of both AKT at Thr308 and GSK3β at Ser9 suggesting that IPMK mediates activation of AKT across cell types (Figure 1G).

**Figure 1.**
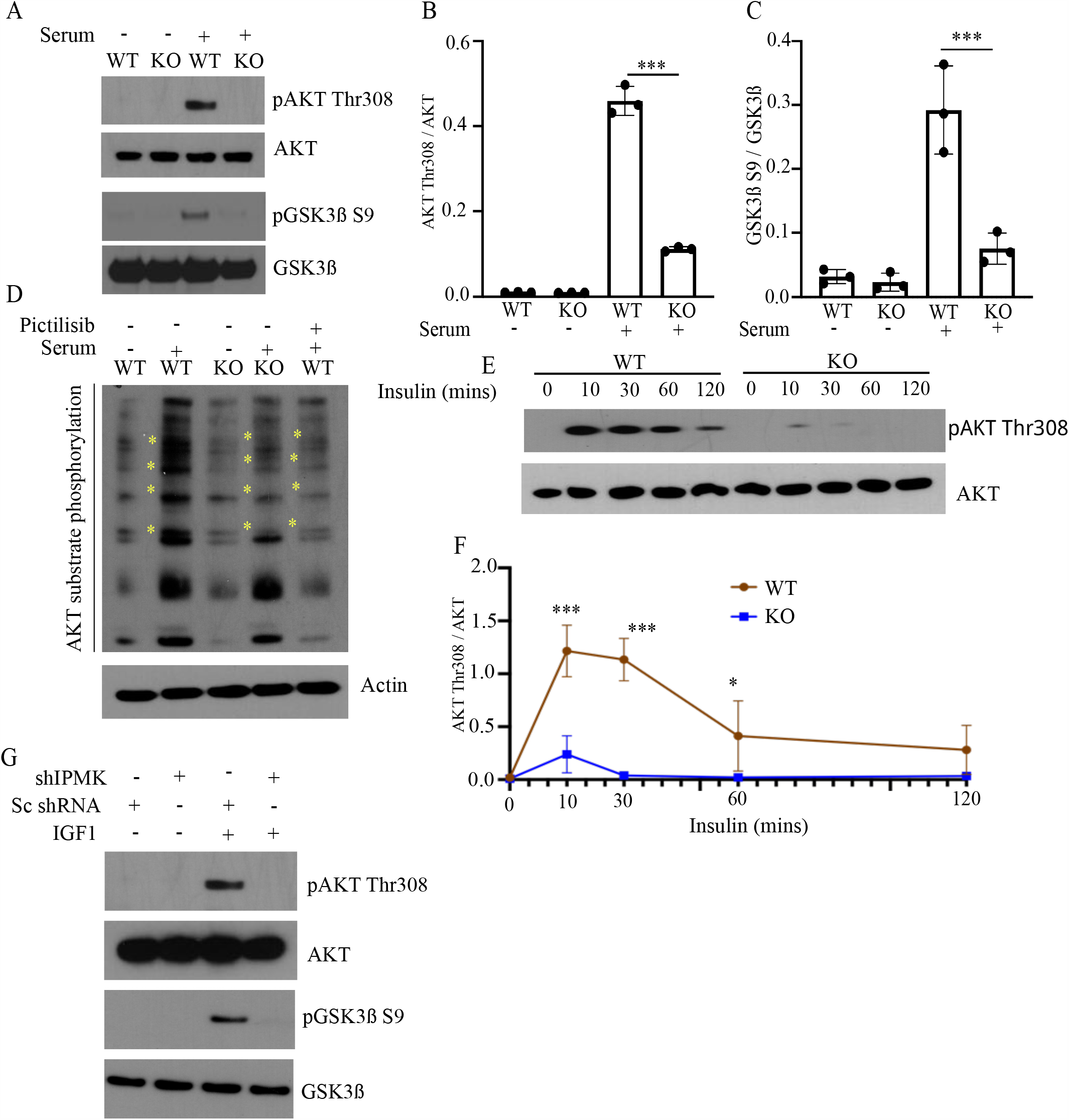
IPMK mediates AKT activation. (**A-C**) Western blot (**A**) and densitometric analysis (**B-C**) of AKT (Thr308) and GSK3β (Ser9) phosphorylation in serum-starved WT and IPMK KO MEFs with or without serum stimulation for 5 min. n = 3. ***p < 0.001. Data are graphed as mean ± SD. (**D**) Western blot of phosphorylated AKT substrates in WT and IPMK KO MEFs treated with serum or pictilisib (100nM, 30 min), n=3. Yellow stars indicates intensity differences. (**E-F**) Western blot (**E**) and densitometric analysis (**F**) of AKT Thr308 phosphorylation in IPMK WT and KO MEFs treated with insulin (10 ng/ml) for the indicated times. n = 3. ***p < 0.001. (**G**) Western blot showing AKT and GSK3β phosphorylation in response to IGF1 treatment (10ng/mL, 10 min) in HEK293 cells treated with control (Sc shRNA) or IPMK (shIPMK) shRNA, n=3.

### PI3-kinase activity of IPMK is required for activation of AKT

IPMK possesses distinct inositol phosphate kinase and PI3-kinase activities ^3^. To ascertain the importance of IPMK’s catalytic activity in regulating AKT’s Thr308 phosphorylation, we rescued IPMK KO MEFs with wild type (wIPMK) and kinase-dead (KSA) IPMK. WT IPMK efficiently rescued AKT Thr308 phosphorylation, whereas KSA IPMK did not (Figure 2 A, B). Thus, the catalytic activity of IPMK is required for AKT Thr308 phosphorylation. IPMK catalyzes the conversion of IP3 to IP4 and IP4 to IP5. Inositol pentakisphosphate-2-kinase (IPPK) catalyzes the conversion of IP5 to IP6 ^3^. Multiple inositol polyphosphate phosphatase 1 (MINPP1) hydrolyzes IP5 and IP6 to IP3 ^15^ (Figure 2 C). To examine the importance of IP5/6 in AKT Thr308 activation, we over-expressed MINPP1 in HEK293 cells, followed by stimulation with insulin-like growth factor 1 (IGF1) for 15 minutes. Cells overexpressing MINPP1 showed comparable AKT Thr308 phosphorylation to empty vector-transfected cells (Figure 2 D). To further confirm that inositol phosphates are dispensable for AKT Thr308 phosphorylation, we transiently knocked down IPPK in HEK293 cells followed by overnight serum starvation and treatment with IGF1. As expected, IGF1 treatment-induced AKT Thr308 phosphorylation in shIPPK cells was comparable to control shRNA cells (Figure 2 E). These experiments confirm that AKT Thr308 activation is mediated by the PI3-kinase activity of IPMK and is independent of inositol phosphates.

**Figure 2.**
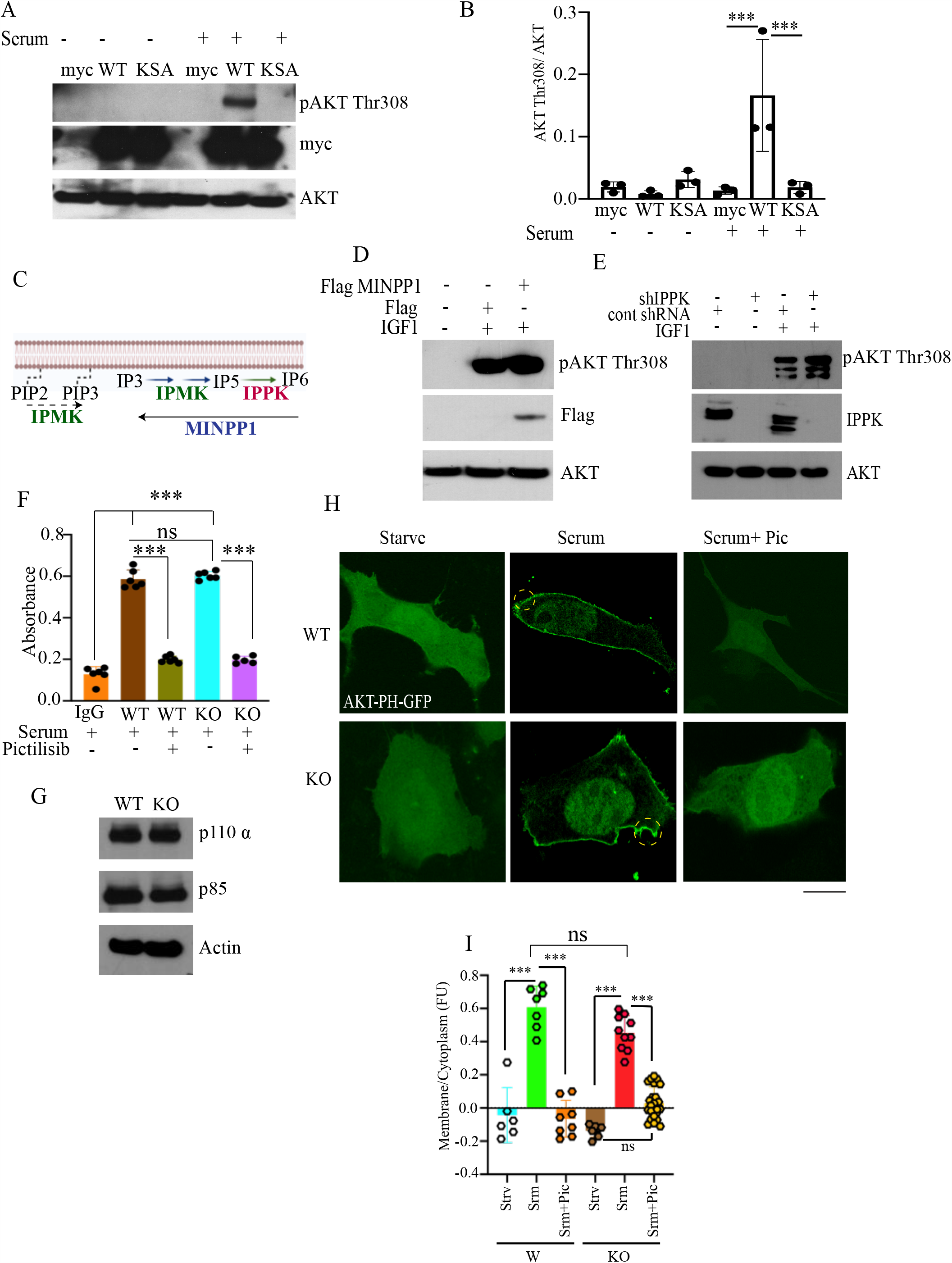
PI3-kinase activity of IPMK is required for activation of AKT. (**A-B**) Western blot (**A**) and densitometric analysis (**B**) of AKT Thr308 phosphorylation in response to serum stimulation in IPMK KO MEFs overexpressing myc-tagged empty vector (myc), WT IPMK (WT), or kinase dead IPMK (KSA) n = 3. ***p < 0.001. (**C**) Cartoon of inositol pathway. (**D**) AKT Thr308 phosphorylation in response to IGF1 treatment in HEK293 cells overexpressing Flag-tagged empty vector (Flag) or MINPP1. n = 3. (**E**). AKT Thr308 phosphorylation in response to IGF1 treatment in HEK293 cells treated with control or IPPK shRNA. n = 3. (**F**) Classical PI3-kinase complex activity was analyzed by pulling down p110α from IPMK WT and KO MEFs treated with serum or pictilisib, n = 3, ***p < 0.001. ns= nonsignificant. (**G**) Western blot of p110α, p85 and actin in IPMK WT and KO MEFs. n = 3 (**H**-**I**) Staining (**H**) and quantification (**I**) of overexpressed AKT-PH-GFP subcellular localization in IPMK WT and KO MEFs in response to serum stimulation and pictilisib (Pic). Membrane-localized AKT-PH-GFP is marked by yellow circle. Scale bar 20μm. Bar diagram shows the relative AKT-PH-GFP membrane/cytoplasm fluorescence ratio. ***p<0.001. Data are graphed as mean ± SD.

We considered the possibility that IPMK influences AKT activation indirectly through actions on classical PI3-kinase activity. To address this possibility, we immunoprecipitated the classical PI3-kinase complex from WT and IPMK KO MEFs and subjected them to an *in vitro* lipid kinase assay. Classical PI3-kinase activity in *Ipmk*-deleted MEFs was comparable to that of wild type cells, as was protein expression of the classical PI3-kinase subunits p110α and p85 (Figure 2 F, G). These results demonstrate that IPMK does not influence classical PI3-kinase activity.

PIP3 promotes AKT membrane localization and subsequent activation ^8^. To investigate the role of IPMK in growth factor-stimulated membrane localization of AKT, we overexpressed the AKT PH domain fused to GFP in WT and IPMK KO MEFs. Serum stimulation efficiently promoted GFP-AKT-PH membrane translocation in WT and KO cells (Figure 2 H, I). By contrast, AKT-PH-GFP membrane localization was prevented by treatment with pictilisib. These immunocytochemical results were further confirmed by cell fractionation and Western blotting of AKT in the cytoplasmic and membrane fractions after serum stimulation (Supplementary Figure 2 A). These results together indicate that IPMK is dispensable for membrane localization of AKT.

### PI3-kinase activity of IPMK mediates PDK1 membrane localization

We wondered whether IPMK directly stimulates the kinase activity of PDK1. To study the influence of IPMK on PDK1 kinase activity, we pulled down PDK1 from WT and KO MEFs and performed an *in vitro* kinase assay using AKT as a substrate. Deletion of *Ipmk* did not impact PDK1-mediated AKT phosphorylation *in vitro* (Figure 3 A). Thus, IPMK has no direct influence on PDK1 kinase activity.

**Figure 3.**
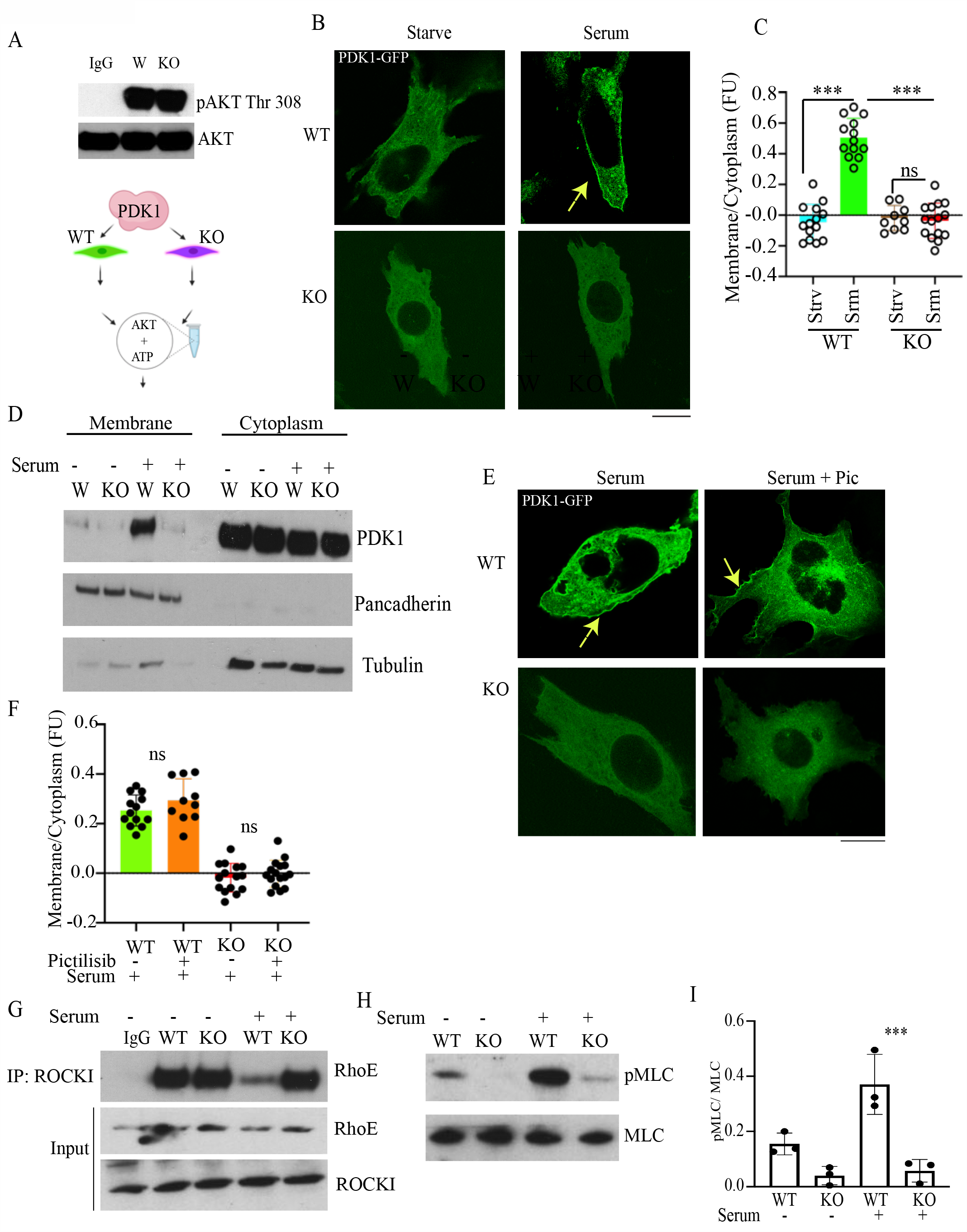
PI3-kinase activity of IPMK mediates PDK1 membrane localization. (**A**) PDK1 *in vitro* kinase assay using AKT as a substrate of PDK1 immunoprecipitated from serum-treated WT and KO MEFs. n=3. (**B-C**) Staining (**B**) and quantification (yellow arrow indicates membrane localized PDK1-GFP) (**C**) of overexpressed PDK1-GFP subcellular localization in IPMK WT and KO MEFs with or without serum stimulation. Membrane-localized PDK1-GFP is marked by yellow arrow. Scale bar 20μm. Bar graph shows the relative PDK1-GFP membrane/cytoplasm fluorescence ratio. ***p<0.001. Data are graphed as mean ± SD. **(D)** Western blot of endogenous PDK1 in cytoplasmic and membrane fractions isolated from WT and IPMK KO MEFs in response to serum stimulation. Pan-cadherin and tubulin blots confirm purity of membrane and cytoplasm fractions, respectively. **(E-F)** Staining (**E**) (yellow arrow indicates membrane localized PDK1-GFP) and quantification (**F**) of PDK1-GFP membrane localization after serum stimulation with or without pictilisib pretreatment (500nM, 30 min). Scale bar 20μm. Bar graph shows the relative PDK1-GFP membrane/cytoplasm fluorescence ratio. ***p<0.001. Data are graphed as mean ± SD. **(G)** ROCK1-RhoE binding in IPMK WT and KO MEFs in response to serum stimulation assessed by immunoprecipitation of endogenous ROCK1 and Western blot of RhoE. (**H-I**) Western blot **(H)** and densitometric analysis (**I**) of MLC phosphorylation (pMLC) in response to serum stimulation in IPMK WT and KO MEFs. n = 3, ***p < 0.001. Data are graphed as mean ± SD.

Serum or growth factor stimulation promotes the generation of PIP3, which binds to the PH domain of PDK1, translocating a fraction of PDK1 protein from the cytoplasm to the cell membrane (20, 23). At the membrane, PDK1 acts as an AKT kinase, phosphorylating AKT at Thr308 (20).

We hypothesized that IPMK-derived PIP3 might influence PDK1 membrane localization. WT and KO MEFs were transiently transfected with full-length GFP-PDK1 followed by overnight serum starvation. As expected, treatment with serum induced export of PDK1 to the cell membrane in WT MEFs, an action that was abolished in KO cells (Figure 3B, C). We confirmed these microscopy data by Western blotting of PDK1 from cytoplasmic and membrane fractions with or without serum stimulation (Figure 3 D).

As observed under confocal microscopy, serum stimulation efficiently triggered AKT membrane localization both in WT and KO MEFs, an action that was abrogated by pictilisib ^11^ (Figure 2F). By contrast, pictilisib had only a marginal effect on PDK1 membrane localization (Figure 3 E, F), suggesting that IPMK is the primary PI3-kinase regulating PDK1 membrane localization.

PDK1 membrane localization is also known to impair ROCK1/RhoE binding, resulting in ROCK1 activation, phosphorylation of myosin light chain (MLC), and cell migration ^12, 13^. Previous studies ^12^ showed that PIP3 promotes localization of PDK1 to the cell membrane, where it dissociates the ROCK1-inhibiting RhoE from ROCK1, effects modestly impacted by classical PI3-kinase inhibitors. We examined the influence of IPMK upon ROCK1/RhoE binding by immunoprecipitation of ROCK1. RhoE/ROCK1 interactions were equal under serum starvation conditions in WT and KO MEFs. Stimulation with serum for 5 minutes markedly depleted ROCK1 binding to RhoE in WT cells, whereas KO cells maintained the ROCK1/RhoE complex at levels comparable to serum starvation (Figure 3G). Myosin light chain (MLC) is a downstream target of ROCK1 and is phosphorylated by the ROCK1-RhoE complex ^13^. ROCK1-mediated activation of MLC promotes cell migration by modulating actin polymerization ^12, 13^. Deletion of IPMK markedly reduced MLC phosphorylation (Figure 3H, I) and cell migration (Supplementary Figure 3A, B), confirming that IPMK promotes PDK1 membrane localization and its downstream signaling events.

### IPMK is essential for intestinal integrity

To examine the influence of IPMK on AKT Thr308 phosphorylation and cell migration *in vivo*, we first examined the localization of IPMK in diverse organs. IPMK expression, assessed by immunohistochemistry (IHC), was highest in the intestinal ileum, while the spleen and skeletal muscle were also enriched (Supplemental Figure 5A). IPMK was abundant in the white pulp of the spleen, whereas it was homogeneously distributed in skeletal muscle. Within cells, IPMK was concentrated both in the nucleus and cytoplasm in the intestine, muscle, and skin hair follicles. IPMK was primarily localized to nuclei of the liver, exocrine glands of the pancreas, cardiac muscle, lungs, and kidney capsule. IPMK expression was highest in the basal region of intestinal ileal crypts (Supplemental Figure 4A).

**Figure 4.**
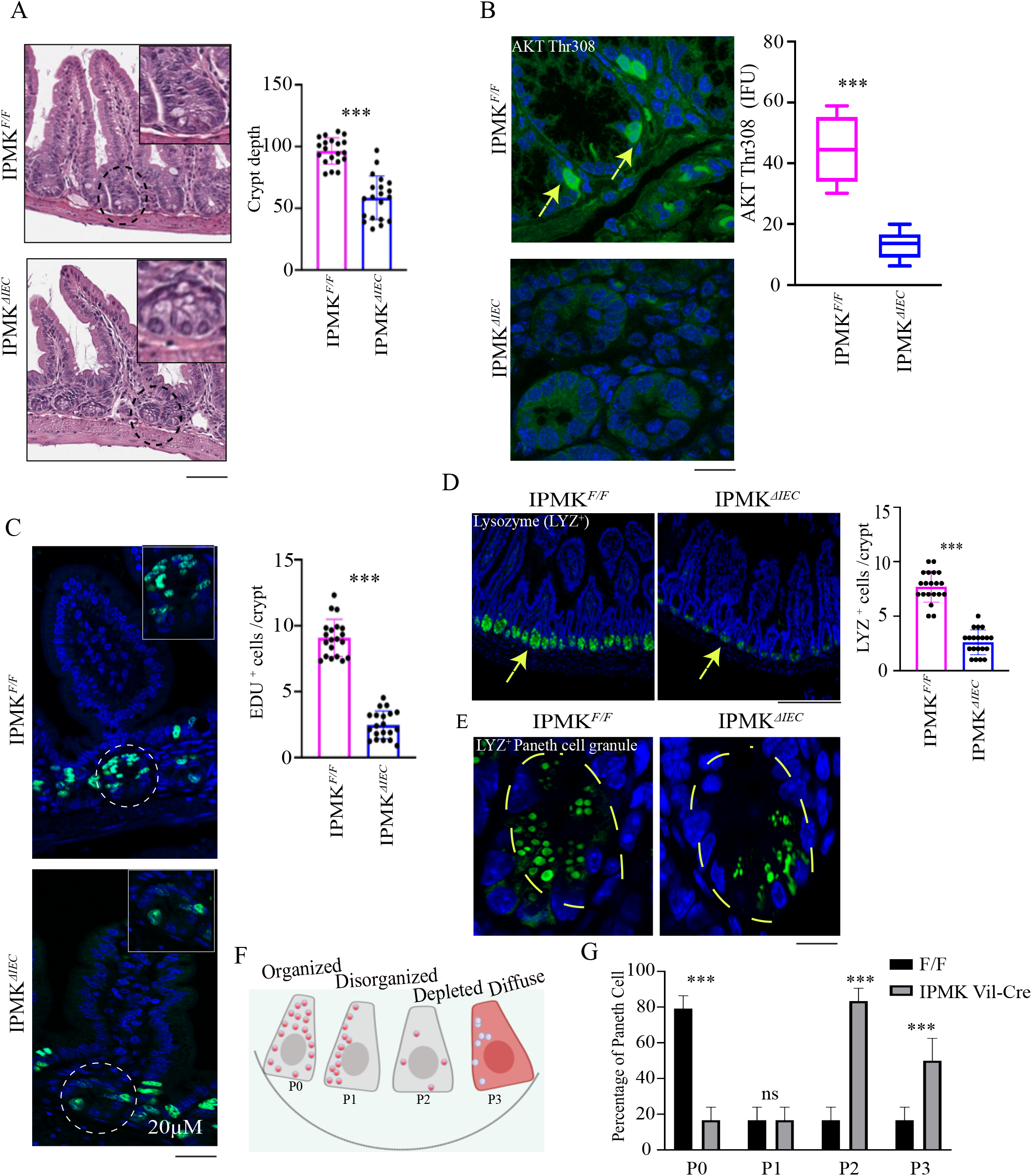
IPMK is essential for intestinal integrity. (**A**) H&E staining of ileum sections from *Ipmk*^*F/F*^ and *Ipmk*^*ΔIEC*^ mice. Graph depicts crypt depth. n=4, ***p < 0.001. Data are graphed as mean ± SD. **(B)** AKT Thr308 staining (green) of ileum sections from *Ipmk*^*F/F*^ and *Ipmk*^*ΔIEC*^ mice, n=4. ***p < 0.001. Data are graphed as mean ± SD. DAPI (blue) used for nuclear staining. **(C)** EdU (green) staining in intestinal sections from *Ipmk*^*F/F*^ and *Ipmk*^*ΔIEC*^ mice intraperitonially injected with EdU 2h prior to sacrifice. DAPI (blue) used for nuclear staining. Number of EdU-positive cells were counted n=4. ***p < 0.001. Data are graphed as mean ± SD. **(D, E)** Paneth cell granules identified by lysozyme (LYZ) staining (green) in ileum sections from *Ipmk*^*F/F*^ and *Ipmk*^*ΔIEC*^ mice, n=4. ***p < 0.001. Data are graphed as mean ± SD. DAPI (blue) used for nuclear staining. **(F-G)** Cartoon (**F**) and quantification (**G**) of lysosomal granule distribution in Paneth cells in *Ipmk*^*F/F*^ and *Ipmk*^*ΔIEC*^ ileum sections. n=5. ***p < 0.001. Data are graphed as mean ± SD. DAPI (blue) used for nuclear staining.

**Figure 5.**
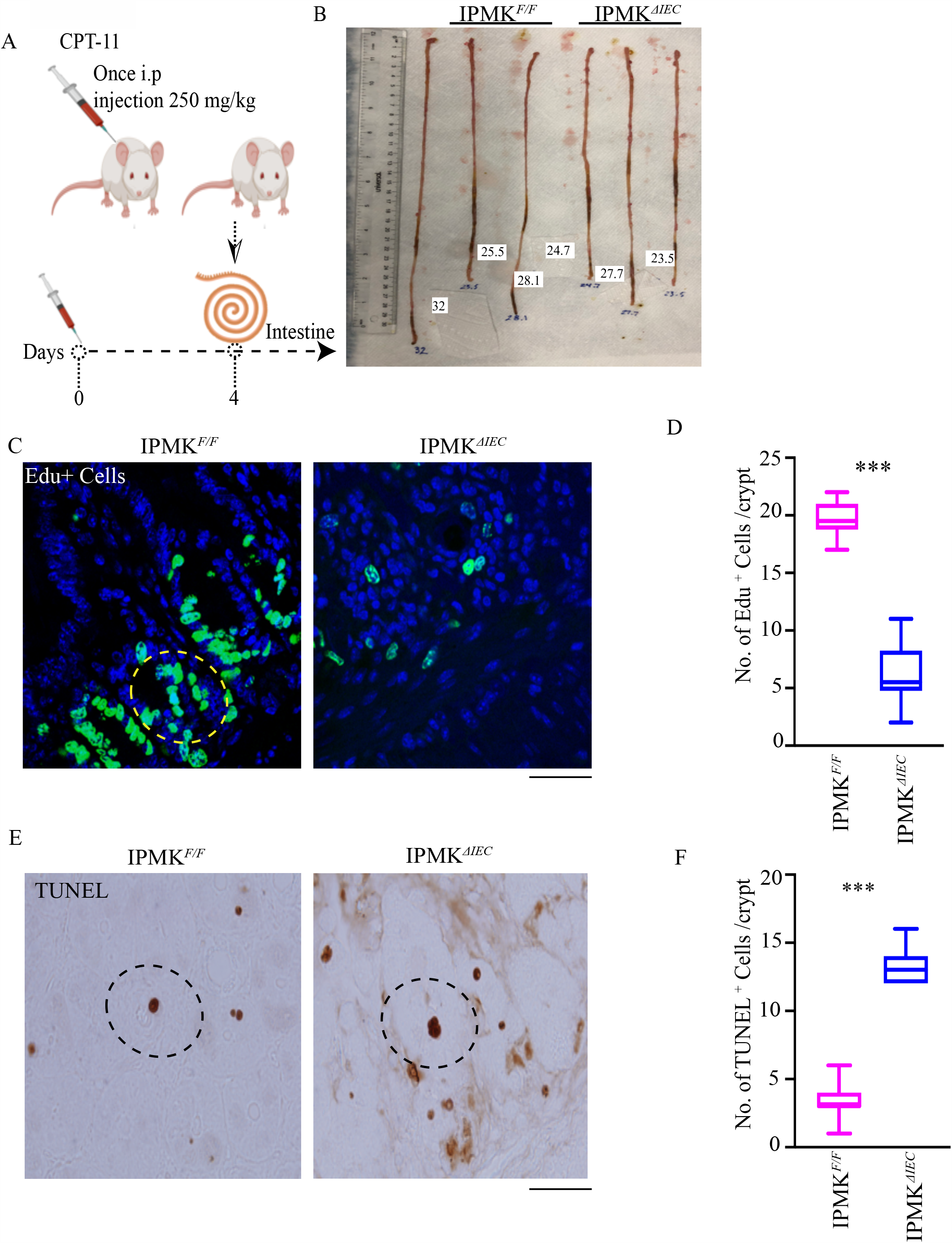
IPMK enhances intestinal epithelial repair. (**A**) Schematic of CPT-11 treatment plan. (**B**) Length of small and large intestine dissected from CPT-11-treated *Ipmk*^*F/F*^ and *Ipmk*^*ΔIEC*^ mice. (n = 3). (**C**) EdU (green) and DAPI (blue) staining 2h after injection in CPT-11-treated intestines. Scale bar 20μm, (yellow circle speify Edu positive cell) (**D**) Quantitation of EdU-positive cells. n=4, ***p < 0.001. Data are graphed as mean ± SD. (**E**) TUNEL-positive cells (dark brown) after CPT-11 treatment in *Ipmk*^*F/F*^ and *Ipmk*^*ΔIEC*^ ileum. Scale bar 20 μm, **(F)** Quantitation of TUNEL-positive cells. n=4, ***p < 0.001. Data are graphed as mean ± SD.

To investigate a potential role for IPMK in activation of AKT and intestinal homeostasis, we crossed *Ipmk* floxed mice (*Ipmk*^*F/F*^) with *Villin-Cre* mice to generate a mouse line lacking *Ipmk* selectively in intestinal epithelial cells (IECs) (*Ipmk*^*ΔIEC*^) (Supplemental Figure 4B). *Ipmk*^*ΔIEC*^ mice were born at the expected Mendelian ratios and developed normally. IEC-specific deletion of *Ipmk* in 8-week-old mice resulted in decreased crypt depth in the ileum (Figure 4A). Cell proliferation in the intestinal epithelium, one of the most proliferative tissues in mammals, requires activation of AKT ^16^. *Ipmk*^*F/F*^ mice showed prominent AKT Thr308 staining, which was markedly diminished in *Ipmk*^*ΔIEC*^ mice (Figure 4 B), demonstrating that IPMK is a physiological regulator of AKT.

AKT is a master regulator of cell proliferation and can play this role through multiple mechanisms. In IECs, phosphorylation of β-catenin at Ser552 by AKT elicits its translocation to the nucleus, where β-catenin acts as a transcriptional coactivator to stimulate cellular proliferation ^17^. Nuclear localization of β-catenin Ser 522 was notably reduced in *Ipmk*^*ΔIEC*^ mice at the intestinal crypt base (Supplemental Figure 4C).

Next, we examined the importance of IPMK in IEC proliferation under basal conditions. *Ipmk*-deleted IECs showed a marked loss of incorporation of the thymidine analog 5-ethynyl-2’-deoxyuridine (EdU), depicting that IPMK is critical for IEC proliferation (Figure 4C). Loss of intestinal regeneration is associated with enhanced cell death ^18^. We observed marked cell death in IECs of *Ipmk*^*ΔIEC*^ mice, whereas *Ipmk*^*F/F*^ mice showed negligible cell death (Supplemental Figure 4D).

Activation of AKT is essential for the homeostasis of Paneth cells that support intestinal regeneration ^19^. Intestinal deletion of IPMK elicited a 50% decrease of Paneth cell number (Figure 4 D). Our examination of periodic acid-Schiff (PAS)/alcian-blue-stained intestinal sections of *Ipmk*-deleted mice revealed pronounced abnormalities in Paneth cells, including aberrant, disorganized granules, as well as decreased numbers of granules (Supplemental Figure 4E). Histological analysis of these sections from 4 controls and 4 mutant mice revealed a 100% concordance between *Ipmk* genotype and abnormal Paneth cell morphology. We quantified these Paneth granule abnormalities by staining intestinal sections for lysozyme, which is usually packaged efficiently in the granules ^20^. We observed large proportions of *Ipmk*^*ΔIEC*^ Paneth cells with depleted lysozyme staining (Figure 4, E-G).

Paneth cell disorganization plays a role in Crohn’s disease pathology ^21^. GWAS data ^7^ revealed a significant association between *IPMK* variants and risk for IBD. Four SNPs at the *IPMK* locus were significantly associated with IBD. SNPs that have statistically significant associations with IBD include rs1819658, located in an enhancer of the *IPMK* gene (P=1.0 × 10^−6^), and rs2790216, located in intron 1 of the *IPMK* gene (P=8.0 × 10^−9^) (Supplemental Figure 5A). Two additional SNPs were also associated with IBD (not presented in Supplemental Figure 5A): rs2153283, located in intron 4 of *IPMK* (IBD P= 2 × 10^−11^, MAF: 0.40, alleles C/A), and rs1199103, an intergenic variant (IBD, P= 5 × 10^−11^, MAF: 0.40, alleles A/C/G).

Epithelial cells are born at the base of the intestinal crypts and migrate to the tip of the villus where they eventually die (Supplemental Figure 5B). A recent study showed that active cell migration is critical for maintaining intestinal homeostasis ^22^. We examined the influence of IPMK on IEC migration 48 h after injection of 5-ethynyl-2’-deoxyuridine (EdU) a thymidine analog which is incorporated into the DNA of proliferative cells. Intestinal deletion of IPMK strikingly impeded IEC migration from the crypt to the villus tip (Supplemental Figure 5C-D). In summary, loss of IPMK expression in IECs led to impaired epithelial cell proliferation, Paneth cell abnormalities, and diminution of IEC migration to the villus tip.

### IPMK enhances intestinal epithelial repair

AKT promotes IEC proliferation during damage-induced intestinal tissue repair ^19^. To investigate a potential role for IPMK in recovery from chemotherapy-induced acute intestinal injury, we treated *Ipmk*^*F/F*^ and *Ipmk*^*ΔIEC*^ mice with a single dose of CPT-11(irinotecan) (250 mg/kg), an agent currently being explored as a treatment for colorectal cancer (Figure 5A), and intestinal tissue was analyzed 4 days later. *Ipmk*^*ΔIEC*^ mice had markedly reduced small intestine size compared to *Ipmk*^*F/F*^ (Figure 5, B and Supplementary Figure 6 B). Histopathological analysis revealed regeneration of crypts and epithelial layers in *Ipmk*^*F/F*^ mice. By contrast, *Ipmk*^*ΔIEC*^ mice displayed significant impairment in the healing of crypts (Supplementary Figure 6 A). We found extensive depletion of EdU-positive cells in *Ipmk*^*ΔIEC*^ mice relative to *Ipmk*^*F/F*^ mice, indicating that IPMK is a mediator of intestinal repair and regeneration (Figure 5C, D). Loss of regeneration in *Ipmk*^*ΔIEC*^ mice was associated with a pronounced increase in apoptosis, as evident from TUNEL staining (Figure 5 E, F). F4/80 staining showed modest invasion of inflammatory cells in *Ipmk*^*F/F*^ mice, demonstrating recovery. *Ipmk*-deleted tissue displayed many inflammatory cells in the intestine, again demonstrating impaired tissue repair and inflammation (Supplemental Figure 6 C). Taken together, these data demonstrate that IPMK plays a critical role in regenerative responses to intestinal toxins.

**Figure 6.**
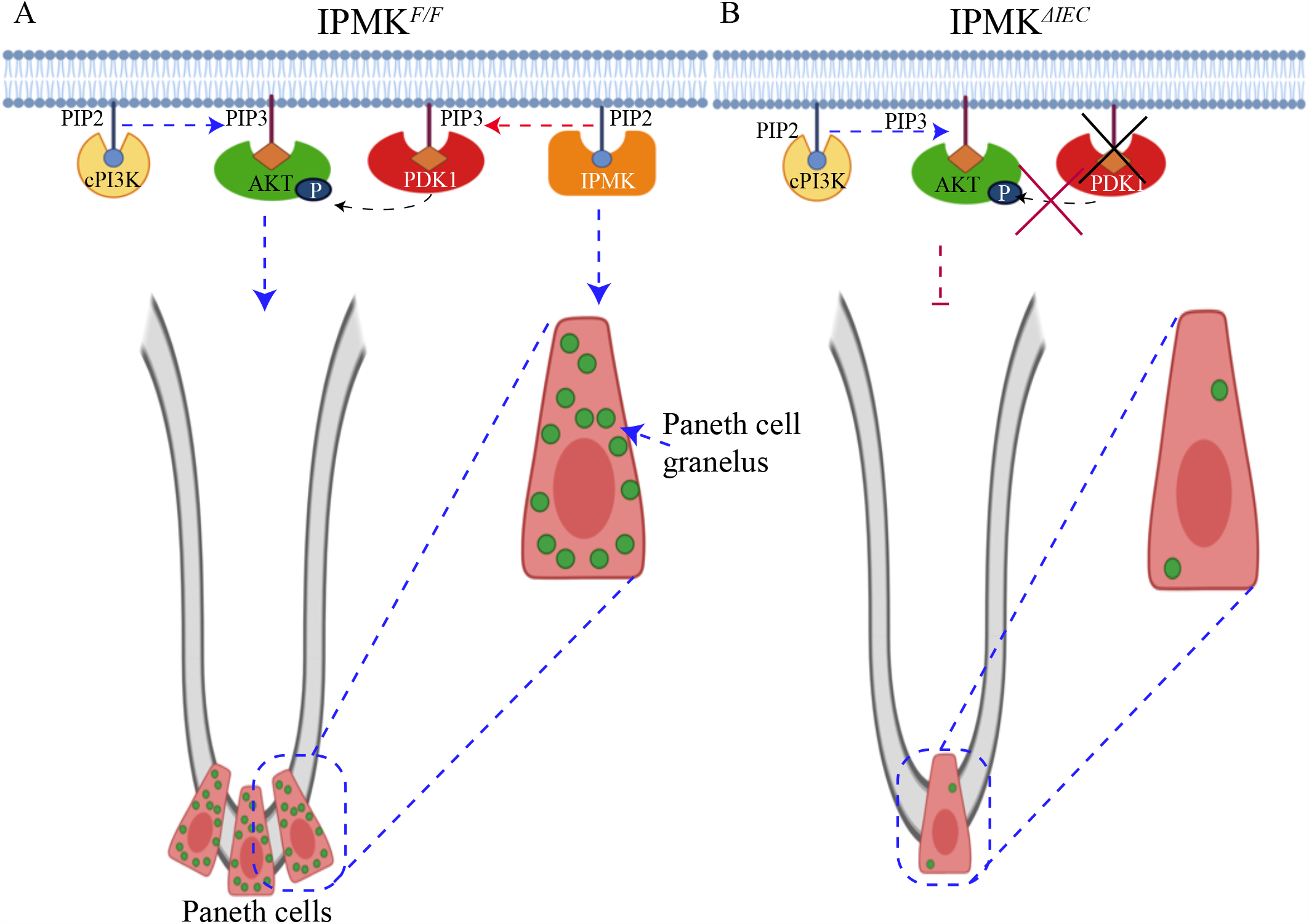
Schematic diagram depicting role of IPMK in PDK1-mediated AKT activation and intestinal homeostasis. (**A**) Under normal conditions, IPMK promotes PDK1 membrane localization and PDK1-dependent AKT phosphorylation. The PI3-kinase activity of IPMK is, however, expendable for AKT membrane localization, which exclusively depends on classical PI3-kinase. PDK1-dependent activation of AKT maintains Paneth cell homeostasis. (**B**) Deletion of IPMK (*Ipmk*^*ΔIEC*^) impairs PDK1 membrane localization and PDK1-dependent AKT phosphorylation, rendering AKT inactive. Loss of function of AKT impairs intestinal proliferation and Paneth cell function in the crypt (e.g., loss of Paneth cell number and dysregulation of intracellular granules).

## Discussion

We previously established IPMK as a novel mammalian PI3-kinase ^3^. However, the physiological implications of IPMK’s PI3-kinase activity have remained obscure. Here we show that the PI3-kinase activity of IPMK is critical for PDK1 membrane localization, which promotes activation of AKT and ROCK1 to promote intestinal epithelial cell migration, proliferation, and regeneration.

It is well accepted that classical PI3-kinase generates PIP3 at the plasma membrane, an action required for AKT membrane localization and PDK1-/mTORC2-mediated AKT phosphorylation. We found that deletion of IPMK abolished AKT phosphorylation at both Thr308 and Ser473 by around 90% (Figure 1 A-B and Supplementary Figure 1 A-B), a deficit which could be rescued by WT IPMK, but not by an IPMK mutant lacking kinase activity (Fig 2 A-E). However, the PI3-kinase activity of classical PI3-kinase (p110/p85) remained intact after IPMK deletion (Figure 2F). Accordingly, we explored mechanisms of IPMK-mediated AKT activation. Confocal imaging and cell fractionation studies demonstrated that AKT membrane localization was independent of IPMK and mediated by classical PI3-kinase (Figure 2 H-I). Previously, we showed ^14^ that deletion of IPMK impaired mTORC2 activation and thus diminished AKT Ser473 phosphorylation (Supplementary Figure 1A, B). We wondered whether IPMK-derived PIP3 controls the kinases that phosphorylate AKT. PDK1 is the first discovered AKT kinase and, like AKT, is recruited to the membrane by PIP3 ^23^. The classical PI3-kinase inhibitor wortmannin ^24^ fails to inhibit PDK1 membrane localization ^10^. Similarly, *Pinner et al*., found that PIP3-mediated PDK1 membrane localization helps to dissociate ROCK1 from inhibitory RhoE binding, thus promoting cell migration; this action was likewise mostly unaffected by classical PI3-kinase inhibitors ^12^. We also showed that a classical PI3-kinase inhibitor had no effect on the PI3-kinase activity of IPMK ^3^. Here we found that the PI3-kinase activity of IPMK is critical for PDK1 membrane localization, which was minimally influenced by the classical PI3-kinase inhibitor pictilisib (Figure 3 B-F). Deletion of IPMK also impaired PDK1-mediated ROCK1 activation by preventing the dissociation of ROCK1 from RhoE (Figure 3 G-I) and diminished cell migration (Supplementary Figure 3A, B). However, recruitment of PDK1 to the membrane does not require physical interaction between PDK1 and IPMK (data not shown). This finding establishes IPMK as a critical mammalian PI3-kinase that promotes membrane localization of PDK1, one of the principal kinases that triggers AKT activation. In contrast to conventional wisdom, PDK1 membrane localization is mostly independent of classical PI3-kinase.

Activation of AKT is essential for maintaining the homeostasis of regenerative organs ^25^. The intestine is a highly regenerative organ, and the loss of AKT function in the intestine impairs intestinal epithelial cell regeneration ^19^. Activation of AKT is critical for maintaining homeostasis of the Paneth cell, a specialized intestinal cell type that resides in ileal crypts and secretes antibacterial peptides and growth factors to regulate intestinal bacterial flora and nurture intestinal stem cells ^26^. IPMK is highly enriched in ileal crypts (Supplementary Figure 4A). As expected, intestinal deletion of IPMK markedly reduced PDK1-mediated AKT Thr308 phosphorylation and IEC proliferation both under basal conditions and during recovery from chemotherapy-induced tissue damage (Figure 4B, C and Figure 5). *Ipmk*^*ΔIEC*^ mice showed a striking loss of Paneth cell number accompanied by pronounced abnormalities in the organization of Paneth cell granules (Figure 4B-G). This Paneth cell abnormality has been observed in Crohn’s disease ^27^. Genome-wide association studies previously identified IPMK as a major Crohn’s disease related gene (Supplementary Figure 5 A). Thus, loss of function of IPMK may drive pathology in Crohn’s disease and IBD by impairing AKT-mediated Paneth cell integrity.

In summary, IPMK is a novel PI3-kinase for PDK1 which activates AKT and promotes ROCK1-mediated cell migration. PDK1 membrane localization and PDK1-dependent AKT phosphorylation is critically regulated by the PI3-kinase activity of IPMK and is independent of classical PI3-kinase. By contrast, the PI3-kinase activity of IPMK is dispensable for AKT membrane localization, which depends exclusively on classical PI3-kinase (Figure 6 A). IPMK-mediated AKT activation promotes intestinal regeneration by maintaining Paneth cell homeostasis. Differences in IPMK activity may therefore contribute to risk for Crohn’s disease, and drugs that target the PI3-kinase activity of IPMK may have potential as novel treatments for Crohn’s disease and related conditions.

## Methods

### Chemicals and reagents

Paraformaldehyde, hydrogen peroxide, antifade mounting medium, FBS, L-glutamine, penicillin/streptomycin, DMEM, PVDF membrane, DMSO cell culture grade, and lipofectamine3000 were purchased from Thermo Fisher Scientific. Irinotecan hydrochloride (CPT-11) was from Sigma (I1406-50MG). ImmPACT® DAB Peroxidase (HRP) Substrate was from Vector Biolab. DyLight 594 Streptavidin was from Vector Biolab (SA-5594). Antigen Retrieval Reagent-Universal was from R&D systems. EZview™ Red Protein G Affinity Gel was from Sigma Aldrich. Click-iT™ EdU Cell Proliferation Kit for Imaging, Alexa Fluor™ 488 was from ThermoFisher. VECTASHIELD® Antifade Mounting Medium and 2.5% Normal Goat Serum Blocking Solution were from Vector Biolab. Citrate Buffer, pH 6.0 was from Sigma Aldrich.

### Antibodies and recombinant proteins

Antibodies against β-Actin (cat no. 4967, 1:3000 dilution), phospho-myosin light chain 2 (Thr18/Ser19) (3674, 1:1000), RhoE (3664, 1:1000), and phospho-AKT (Ser473) (4060, 1:2000, for Western blot) were from Cell Signaling Technology. Antibodies against phospho-AKT (Thr308) (GTX79150, IHC 1:100, WB 1:2000). Lysozyme antibody (ab108508, 1:100) was from Abcam. Phospho-β-catenin-S552 (AP0579, 1:100) was from ABclonal. Myosin Light Chain 2 antibody (10906-1-AP, 1:1000) was from Proteintech. ROCK1 antibody (A300-457A, 1:1000) was from Bethyl Laboratories. Goat anti-Rabbit, Alexa Fluor 488 antibody (A-11008, 1:1000) was from Invitrogen.

## Cell culture

Mouse embryonic fibroblast (MEF) and human embryonic kidney (HEK) cells were cultured in Dulbecco’s modified Eagles medium (DMEM) supplemented with 10% FBS, 2 mM L-glutamine, 100 U/mL penicillin, and 100 mg/mL streptomycin. IPMK WT and KO MEF cells were developed in our lab as previously described.^3^

### Generation of intestine-specific IPMK KO mice

*Ipmk*^*F/F*^ mice were generated as previously described (22). *Ipmk*^*F/F*^ mice were crossed with C57BL/6J mice carrying Cre expressed under the control of the murine villin promoter as described previously (34) to create intestinal-specific *Ipmk* knockout mice (*Ipmk*^*ΔIEC*^). Homozygous *Ipmk*^*F/F*^ mice were crossed with the *Ipmk*^*ΔIEC*^ mice, which mediate excision of floxed alleles in the intestine. All mice were maintained on a C57BL/6J background and were 7th generation backcrossed. C57BL/6J mice were used for Enteroid isolation for inhibitor treatment as well as for DSS and CPT-11 treatments. Mice were housed in a 12-hour light/12-hour dark cycle at 22°C and fed standard rodent chow. All research involving mice was approved by the Johns Hopkins Animal Care and Use Committee.

### Plasmids and transfection

AKT-PH-GFP and GFP-PDK1 plasmids were from Addgene. pMX-myc, pMX-wIPMK and pMX-IPMK-KSA were generated in-lab.

### Retroviral Transfection and Generation of Stably Transfected Cells

Target vector DNA retroviral constructs were transiently transfected into a Platinum-E (Plat-E) retrovirus packaging cell line for 48 hours by using Lipofectamine 3000 transfection reagent. High-titer viral stocks were harvested by passing the supernatant through a 0.45 μm filter. For infection, MEFs were incubated with the viral supernatant in the presence of polybrene (8μg/mL) for 48 hours. Stably infected MEFs were selected by treatment with blasticidin (4 μg/mL) for 1-2 weeks. Selected stable cell lines were continuously maintained in medium containing blasticidin (4 μg/mL).

### Transient Transfection

MEF cells were transfected with Lipofectamine 3000 (ThermoFisher) per manufacturer protocol.

### AKT activation in cell culture

To evaluate AKT phosphorylation in MEF cells, 5×10^6^ cells were seeded in 10 cm plates and left to attach to the plate for 3 hours. Cells were then washed with room temperature phosphate-buffered saline (PBS) followed by incubation of cells in serum-free DMEM supplemented with 2 mM L-glutamine, 100 U/mL penicillin, and 100 mg/mL streptomycin. After 24 hours of serum starvation, serum-free media was replaced with complete DMEM (10% FBS) for 5 minutes to promote AKT membrane localization. Cells given the PI3K inhibitor GDC0941/pictilisib were pretreated for 30 minutes prior to complete DMEM administration with GDC0941 (100 nM) in serum-free media. After 5 minutes of serum stimulation, cells were quenched with cold PBS and lysed as previously described.

### Cytoplasm and cell membrane isolation

Cytoplasm and membrane fractions were obtained using the ProteoExtract® Subcellular Proteome Extraction Kit (Millipore-Sigma] following manufacturer protocol.

### Immunoprecipitation

Immunoprecipitation of endogenous ROCK-I was performed with 1 mg of protein lysates in lysis buffer (150mMNaCl, 0.5% CHAPS, 0.1% Triton, 0.1% BSA, 1mM EDTA, protease inhibitors, and phosphatase inhibitors). Protein lysate was incubated for 2 hrs. at 4*C with ROCK1 antibody (Bethyl) followed by EZview Protein G beads (Sigma) for 1 h. Beads were pelleted at 1000xg and washed with lysis buffer 3 times for 5 minutes each on a rocking platform at 4*C, and SDS sample buffer loading dye was added. Immunoprecipitated samples were resolved by polyacrylamide gel electrophoresis and binding of RhoE was observed by Western blot.

### In vitro PDK1 kinase activity assay

To perform In vitro PDK1 kinase assay 3 µg of PDK1 antibody was added to 1.5 mg of lysates and incubated with rotation for 2 hours at 4°C. 15 µL of protein G beads was added and the incubation continued for an additional hour. PDK1 immunoprecipitates were washed four times with the same lysis buffer and twice with the mTORC2 kinase buffer (25 mM HEPES [pH 7.5], 100 mM potassium acetate, 1 mM MgCl2). Kinase assays were performed for 20 min at 37 °C in a final volume of 15 microliters of kinase buffer containing 500 μM ATP and 500 ng inactive AKT1/PKB1 (Millipore) as a substrate. Reaction was stopped by resuspending beads in SDS-containing sample buffer and boiling samples for 5 minutes followed by western blot of phosphor-AKT Thr308.

### AKT and PDK1 membrane localization

5×10^5^ MEF cells were seeded on 35 mm glass-bottom cell culture dishes and left to proliferate for 3 hours. Cells were then transfected with 500 ng AKT-PH-GFP using Lipofectamine 3000 following manufacturer’s recommendations. 24 hours after transfection, cells were washed with PBS and deprived of serum by exchanging complete DMEM with FBS-free media as previously described. After 24 hours of serum starvation, serum-free media was replaced with complete DMEM (10% FBS). Cells given the PI3K inhibitor GDC0941/pictilisib were treated as described above. After 5 minutes of serum stimulation, cells were quenched with cold PBS and fixed using chilled 4% paraformaldehyde. Cells were stained with DAPI (1 μg/mL) in PBS. After staining, coverslips were mounted with antifade (Vectashield) mounting media. Cells were imaged via confocal microscopy; images were exported as .czi files to ImageJ (NIH). For each image, well-defined and intact portions of the membrane were traced using the “segmented line” function with spline-fit, setting the line thickness such that the selection covered 2.5 μm across the membrane. This was followed by transformation using the “straighten” function to generate a uniform, straight section of membrane. This image was rotated such that the left of the image was the intracellular space and the right side was the extracellular space, with the edge of the membrane lying in the middle of the image. A plot profile was generated using this image, yielding a trace of the gray value across the membrane, progressing intra- to extracellular down the x-axis. The data produced by this plot was exported to Excel. As the transfection efficiency differed slightly between cells, data from each image was normalized from 0 to 1, with 0 being the extracellular space and 1 being the maximum gray value. Using this data, the difference between cytoplasm and membrane fluorescence was calculated.

### Western Blot

Cell lysates were first prepared using lysis buffer (50mM Tris-HCl, 150mM NaCl, 1mM EDTA, 1% Triton X-100 in PBS, phosphatase inhibitors, protease inhibitors). Samples were centrifuged at 16,200xg for 10 min, followed by total protein quantification. Proteins were resolved by SDS-polyacrylamide gel electrophoresis and transferred to PVDF membranes. Membranes were incubated in 1:1000 dilution of primary antibody in 20 mM Tris-HCl (pH 7.4), 150 mM NaCl, and 0.02% Tween 20 (Tris-buffered saline/Tween 20, TBST) with 3% BSA overnight at 4°C on a rocking platform. The membranes were washed 3 times with Tris-buffered saline/Tween-20, then incubated with HRP-conjugated secondary antibody (GE Health Care) diluted 1:5000 in TBST with 3% BSA, and the bands visualized by chemiluminescence (ThermoFisher). The blots shown in the figures were representative replicates selected from at least 3 experiments. Densitometric analysis was performed using ImageJ software.

### Immunohistochemistry, imuunofluorescence and DAB stain

Mice were sacrificed via CO_2_. Immediately following euthanasia, the intestine was removed and flushed with cold PBS. The intestine was subsequently opened lengthwise and washed in cold PBS, then cut into approximately 3-4 mm fragments for sectioning and paraffinization. Samples were deparaffinized with Histo-Clear (Thermo Fisher) and rehydrated in successive washes of 100%, 90%, and 75% ethanol followed by deionized (DI) water. Sections were unmasked via heat-induced epitope retrieval (HIER) using citrate buffer, followed by blocking with 2.5% goat serum for 1 hour. Primary antibodies were diluted per manufacturer’s recommendation in Triton-X diluted to 1% in PBS, incubating overnight at 4°C in a humidified chamber. The following day, samples were washed with TBST, followed by incubation with fluorescent-tagged secondary antibodies diluted in 1% Triton-X solution for 60 minutes. After washing with TBST, samples were stained with 4′, 6-diamidino-2-phenylindole (DAPI) diluted in PBS (1 µg/mL). Slides were imaged using slide scanner and confocal microscopy.

To identify the influence of IPMK deletion on Paneth cell we used lysozyme to detect Paneth cells. These, along with anti-Beta-catenin Ser 522, anti-IPMK and IgG control antibodies, were all anti-rabbit and targeted with fluorescent secondary Alexa Fluor488. Additionally, F4/80 staining (used to detect inflammatory cells) was detected using Alexa Fluor 488-tagged anti-mouse secondary antibodies. Phospho-AKT Thr308 antibodies were targeted with Alexa Fluor488-tagged anti-rabbit antibodies.

Tissue samples of small intestine, large intestine, lung, kidney, liver, skeletal muscle, cardiac muscle, pancreas, skin, and lymph were sectioned and paraffinized as described above. Endogenous peroxidase activity was first blocked with [0.1%] H_2_O_2_ for 5 minutes. Sections were incubated with IPMK antibody followed by HRP-tagged secondary antibody, then treated with diaminobenzidine (DAB) substrate. Sections were counterstained with hematoxylin. Images were acquired at 40x magnification with an Olympus microscope. Hematoxylin and DAB channels were separated using the ImageJ color deconvolution macro, which utilizes an RGB image deconvolution method developed by Ruifrok and Johnston (36). Subsequent histogram analysis in ImageJ produced signal intensities from 0-255, with a value of 0 = darkest intensity (positive staining) and 255 representing a white, unstained signal. A threshold of 0-230 was applied to eliminate background staining; using this threshold, the MGV of the tissue samples was determined. Using the MGV, the optical density (OD) of each sample was determined using the equation 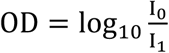, where I_0_ is the intensity of transmitted light (255) and I_1_is the intensity of the DAB-positive area of the sample.

### TUNEL staining

Histologic sections were stained using TumorTACS™ In Situ Apoptosis Detection Kit (Trevigen) following manufacturer protocol.

### Periodic acid Schiff staining

Samples were first deparaffinized with Histo-Clear and rehydrated in successive washes of 100%, 90%, and 75% ethanol, followed by a wash in deionized (DI) water. Slides were stained using the Periodic Acid Schiff Stain Kit (Abcam) following the manufacturer’s protocol.

### EdU treatment and tissue regeneration study

To evaluate DNA synthesis as a marker for epithelial regeneration, mice were given a single administration of 5–ethynyl–2′–deoxyuridine (EdU) by intraperitoneal injection at [200ug/kg] in PBS. Mice were sacrificed 2 hours after administration, with both small and large intestines prepared for histological analysis using Click-iT™ EdU Cell Proliferation Kit for Imaging, Alexa Fluor™ 488 (Thermo Fisher) according to manufacturer protocol.

### Intestinal and MEF Cell Migration

To check cell migration in MEF we performed scratch wound assay. WT and KO MEFs were plated in a glass bottom tissue culture plate with a silicon spacer in the middle. Once cells were adhered they were kept in serum starved medium overnight. Next day the serum free medium was replaced by plus serum medium and spacer was gently taken out. After 7 h the cell migration was captured under light microscope. The path travelled by the cells were analyzed using Fiji2 software.

Intestinal villi length was measured from the base to the apex using segmented line tool in the Image J software. For each villi the length was subdivided into 3 equal sections (base, stem, apex), cells were counted and reported as per these subdivisions. For each subdivision, cell counts are reported as an average, by dividing the number of cells counted per subdivision by the total number of villi (*Ipmk*^*F/F*^ 40, *Ipmk*^*ΔIEC*^ 42).

### CPT-11 treatment and tissue repair study

Mice were given a single administration of CPT-11 (Sigma) by intraperitoneal injection at 250 mg/kg in saline, in a total injection volume of 200 µl. Mice were sacrificed 4 days post-injection as described above, with both small and large intestine taken for analysis.

### Confocal microscopy

Images of adherent cells were acquired with a Zeiss LSM700 single-point, laser scanning confocal microscope. Histologic sections were imaged with both the Zeiss LSM700 as well as an ImageXpress Micro XLS widefield high-content analysis system. Images were analyzed with Zenlite, Imaris, and ImageJ software.

### Statistical analysis

All plots and statistical analysis were performed with Prism 8 (GraphPad) software. Statistical significance was determined by either Student’s t-test (two-tailed) for 2 groups or 1-way ANOVA for multiple groups with similar sample sizes. Error bars represent standard error of the mean, and n indicates number of experimental replicates or the number of animals employed. Differences between groups were considered significant when *p<0.05, **p<0.01, and ***p<0.001.

## ACKNOWLEDGMENTS

We sincerely thank Dr. Gregg Semenza for his critical comments to shape the manuscript. This work was supported by National Institutes of Health grant 5R01MH018501-48.

## AUTHOR CONTRIBUTIONS

PG and SHS conceived and PG designed the experiments. PG, LR, ERS, EA, and YS performed the experiments. MS performed analysis of intestinal cell migration. PG, SAW, MD, and SHS analyzed the data. PG, ERS and MS edited the manuscript. PG and SHS wrote the manuscript. All authors reviewed and commented on the manuscript.

## DISCLOSURES

The authors declare no competing interests.

---

**Supplementary Figure 1.**
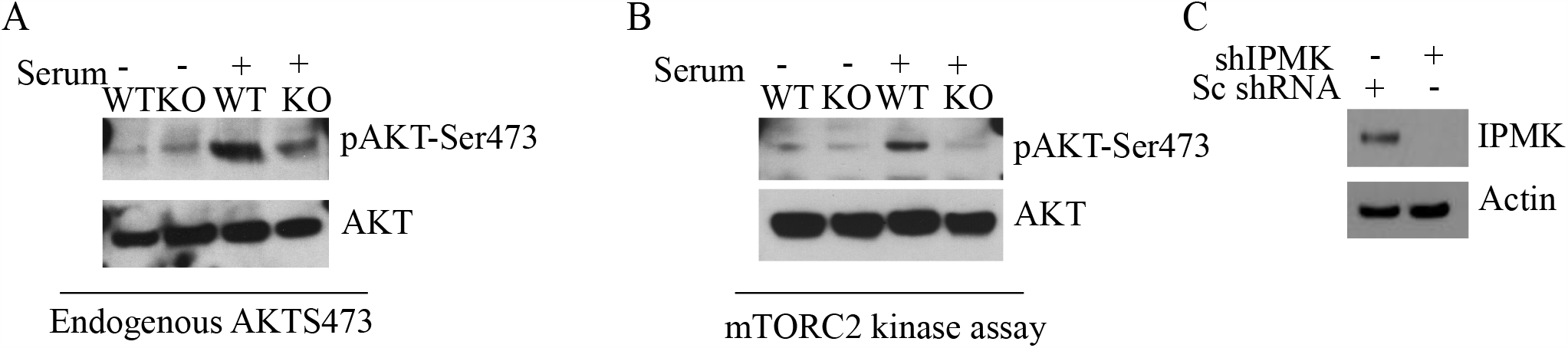
IPMK regulates AKT phosphorlation. (**A**) Western blot of AKT phosphorylation at Ser473 in WT and IPMK KO MEFs serum starved overnight followed by 5 mins of serum treatment. n=4. **(B)** *In vitro* mTORC2 kinase assay in WT and IPMK KO MEFs using AKT as a substrate. mTORC2 complex was immunoprecipitated using rictor antibody. AKT Ser473 phosphorylation was analyzed to analyze mTORC2 kinase activity. n=4. **(C)** Western blot confirming knockdown of IPMK after IPMK shRNA treatment in HEK293 cells.

**Supplementary Figure 2.**
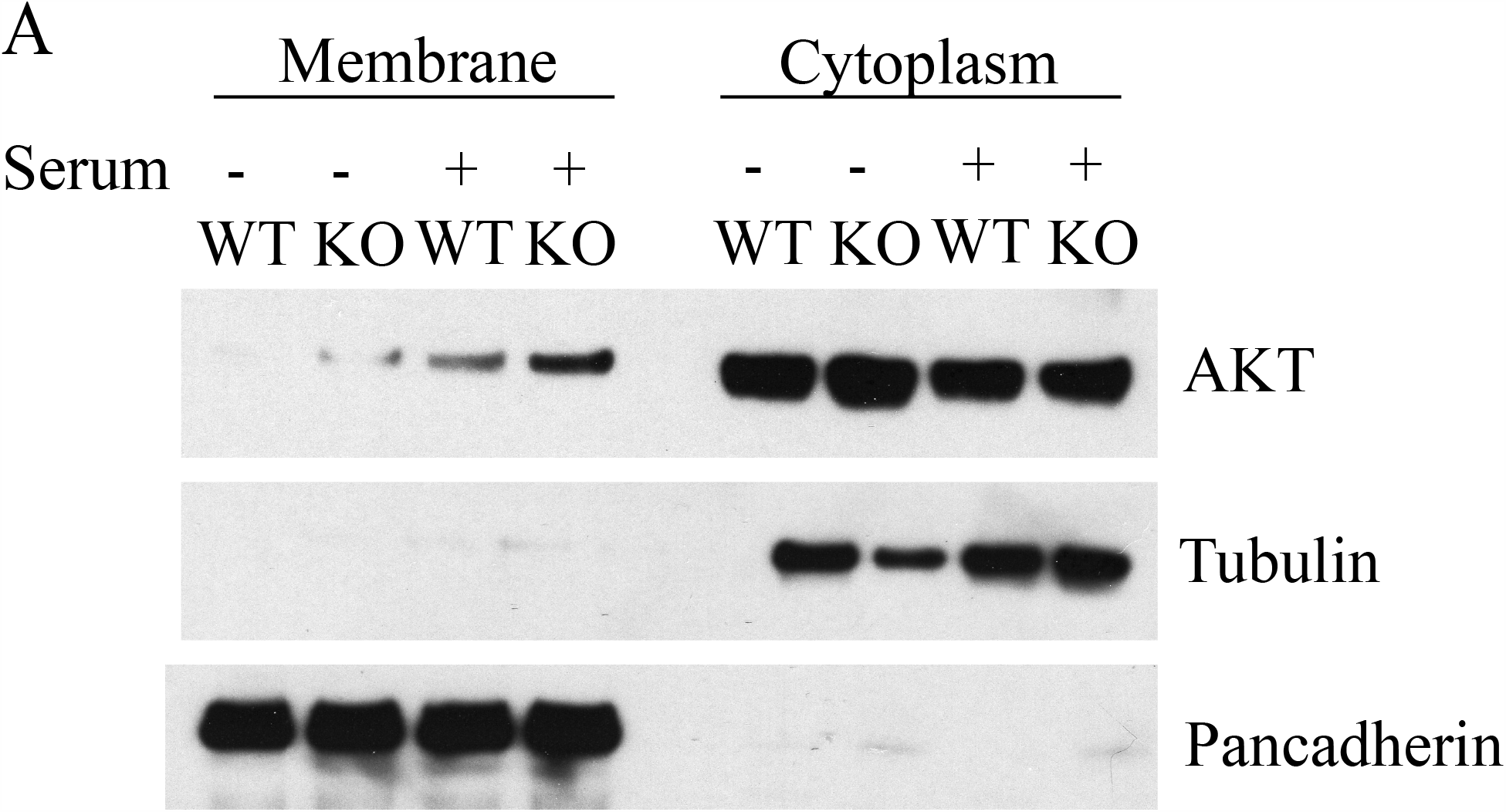
PI3-kinase activity of IPMK is required for activation of AKT. **(A)** Western blot of AKT in membrane and cytoplasmic fractions isolated from WT and IPMK KO MEFs in response to serum stimulation. Pan-cadherin and tubulin blots confirm purity of membrane and cytoplasm fractions, respectively. n=3.

**Supplementary Figure 3.**
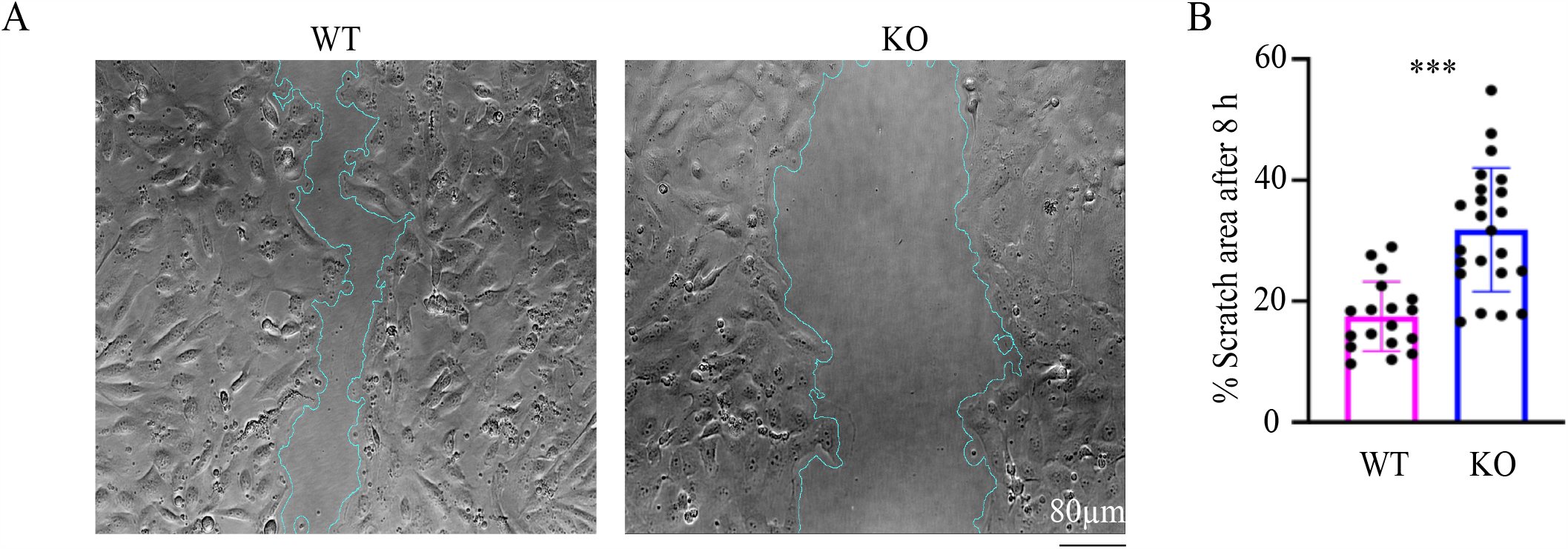
IPMK regulates cell migration. **(A, B)** IPMK WT and KO MEFs were subjected to scratch wound assay to analyze cell migration. Scale bar 80μm. n = 3, ***p < 0.001. Data are graphed as mean ± SD.

**Supplementary Figure 4.**
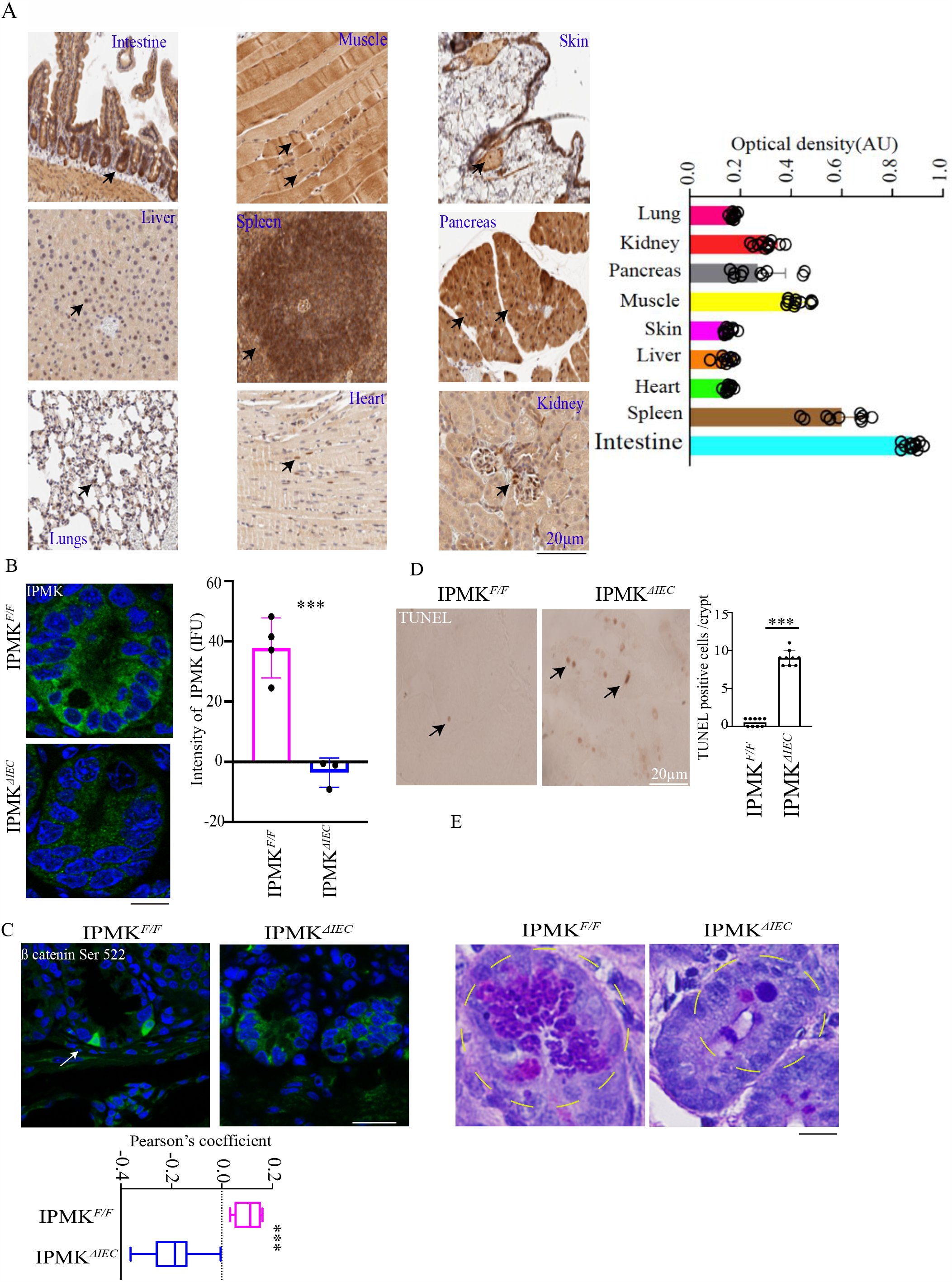
Tissue distribution and intestinal epithelial cell conditional knockout of IPMK. (**A**) IHC study of IPMK expression in different mouse organs (**B**) Staining and quantification of IPMK levels in *Ipmk*^*ΔIEC*^ and *Ipmk* ^F/F^ intestines. Arrow indicates IPMK localization. Values are normalized to fluorescence intensity of sections incubated with a preimmune IgG instead of IPMK antibody. n = 3. **(C)** Staining and quantification of nuclear localization of β-catenin phosphorylation at Ser552 (green) in intestinal sections from *Ipmk*^*F/F*^ and *Ipmk*^*IEC*^ mice. Nuclei are stained with DAPI (blue). White arrows depict phospho-β-catenin (green) nuclear (blue) localization. Scale bar 20μm. n = 3, ***p<0.001. Data are graphed as mean ± SD. (**D**) TUNEL staining and quantification in *Ipmk*^*ΔIEC*^ and *Ipmk* ^F/F^ ileum sections. Scale bar 20μm. ***p<0.001. Data are graphed as mean ± SD. (**E**) Periodic acid–Schiff (PAS) staining of *Ipmk*^*ΔIEC*^and *Ipmk* ^F/F^ ileum sections. Scale bar 20μm. n=4.

**Supplementary Figure 5.**
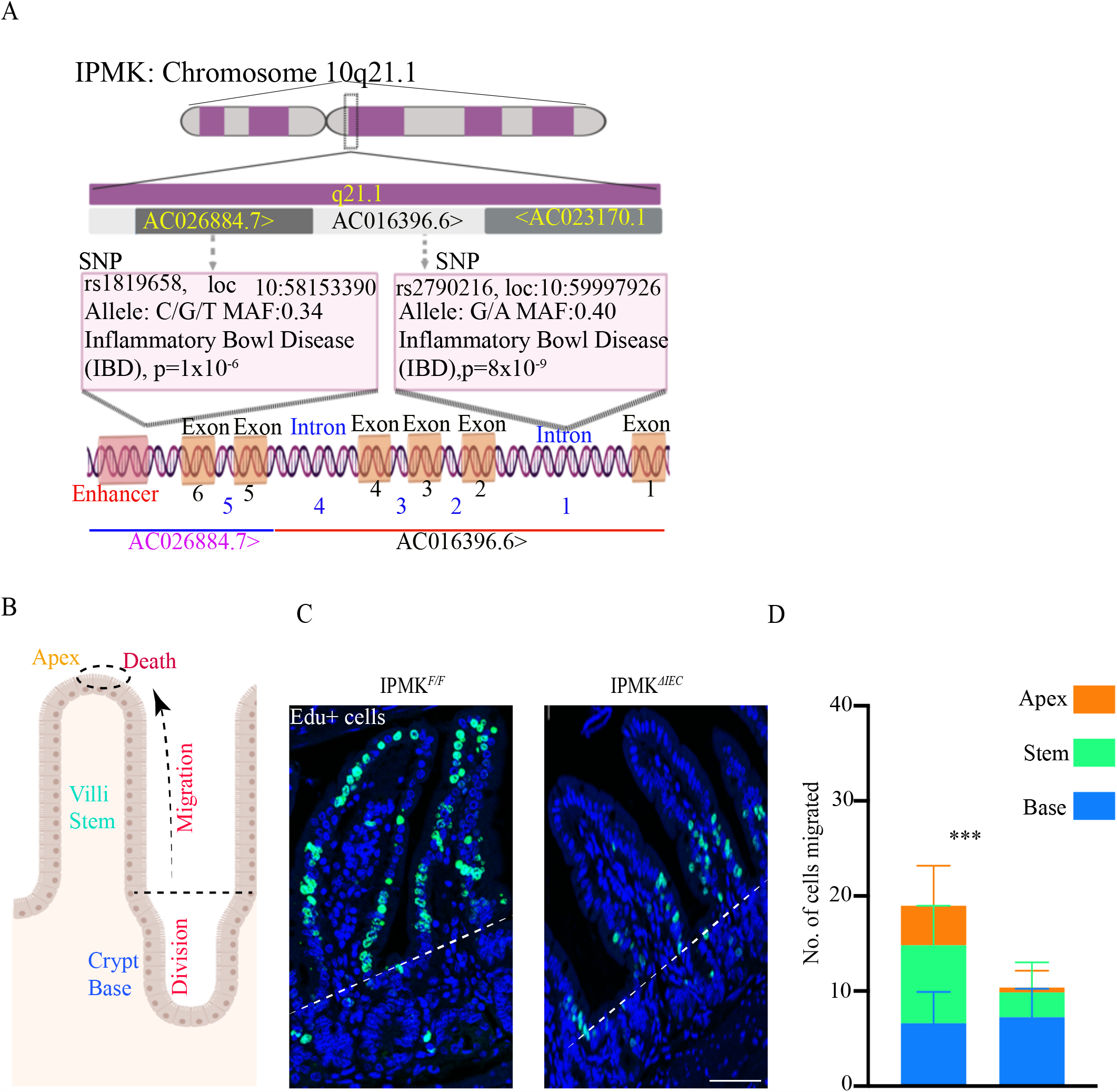
IPMK is related to inflammatory bowel disease and intestinal cell migration. (**A**) Schematic diagram of single nucleotide polymorphisms in *IPMK* implicated in inflammatory bowel disease. (**B**) Cartoon depicting division and migration cycle of intestinal epithelial cells (IECs). (**C-D**) EdU (green) and DAPI (blue) staining, scale bar (**C**) and quantification (**D**) of IEC migration in *Ipmk*^*ΔIEC*^ and *Ipmk* ^F/F^ ileum sections 48 h after intraperitoneal injection of EdU. Scale bar 40μm. n = 3, ***p < 0.001. Data are graphed as mean ± SD.

**Supplementary Figure 6.**
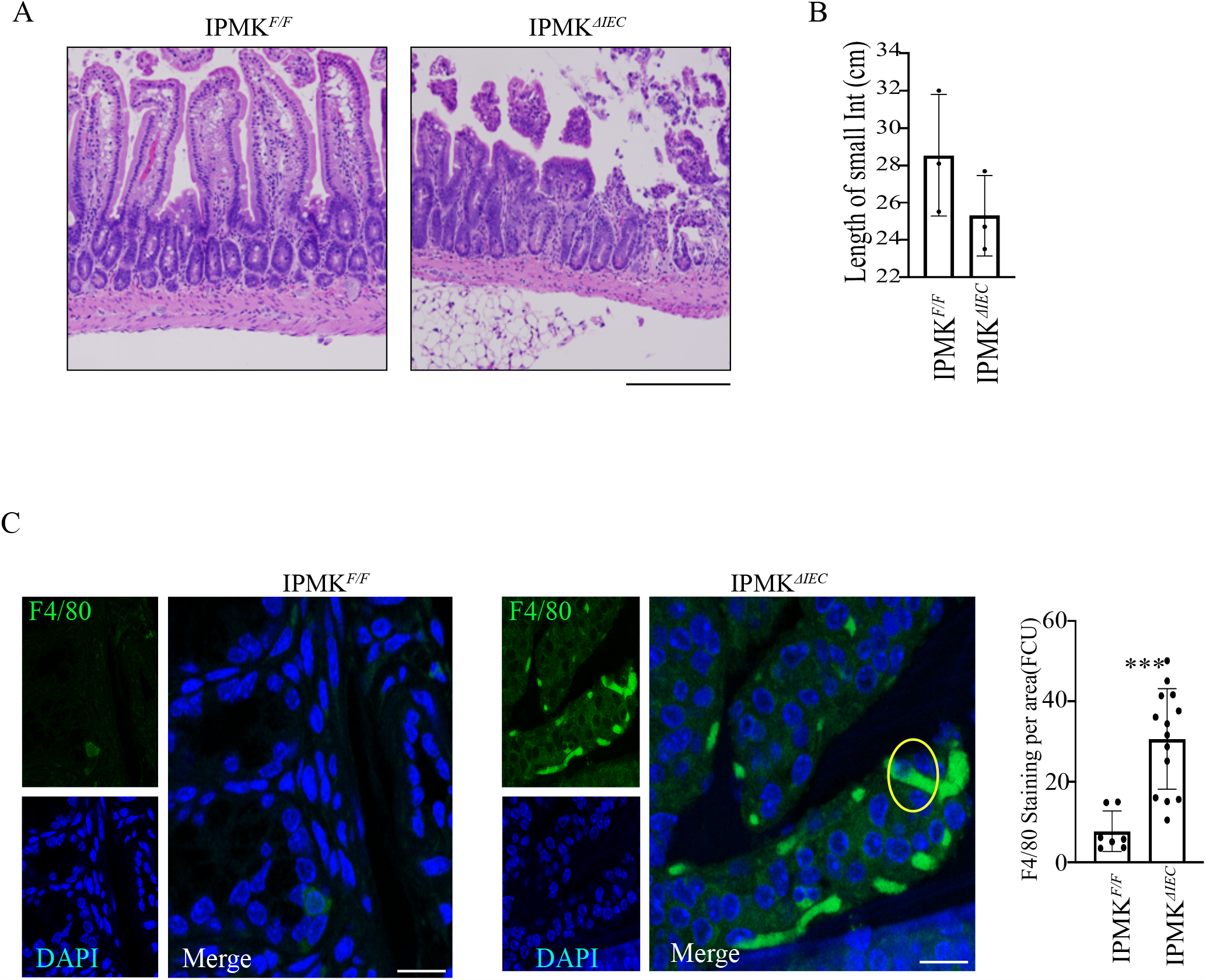
IPMK modulates CPT-11-induced intestinal injury. (**A**) H&E staining of CPT-11-treated ileum sections from *Ipmk*^*ΔIEC*^ and *Ipmk* ^F/F^ mice. (B) Length of ileum after CPT-11 treatment. n=3. (**C**) Staining and quantification of F4/80 (green) in intestinal lamina propria of *Ipmk*^*ΔIEC*^ and *Ipmk* ^F/F^ mice. Scale bar 20μm. n = 3, ***p < 0.001. Data are graphed as mean ± SD.

## Notes

### Competing Interest Statement

The authors have declared no competing interest.

### Summary of Updates

The author names were not right in the web version

